# Insight into vaccine development for Alpha-coronaviruses based on structural and immunological analyses of spike proteins

**DOI:** 10.1101/2020.06.09.141580

**Authors:** Yuejun Shi, Jiale Shi, Limeng Sun, Yubei Tan, Gang Wang, Fenglin Guo, Guangli Hu, Yanan Fu, Zhen F. Fu, Shaobo Xiao, Guiqing Peng

## Abstract

Coronaviruses that infect humans belong to the *Alpha-coronavirus* (including HCoV-229E) and *Beta-coronavirus* (including SARS-CoV and SARS-CoV-2) genera. In particular, SARS-CoV-2 is currently a major threat to public health worldwide. However, no commercial vaccines against the coronaviruses that can infect humans are available. The spike (S) homotrimers bind to their receptors through the receptor-binding domain (RBD), which is believed to be a major target to block viral entry. In this study, we selected *Alpha-coronavirus* (HCoV-229E) and *Beta-coronavirus* (SARS-CoV and SARS-CoV-2) as models. Their RBDs were observed to adopt two different conformational states (lying or standing). Then, structural and immunological analyses were used to explore differences in the immune response with RBDs among these coronaviruses. Our results showed that more RBD-specific antibodies were induced by the S trimer with the RBD in the “standing” state (SARS-CoV and SARS-CoV-2) than the S trimer with the RBD in the “lying” state (HCoV-229E), and the affinity between the RBD-specific antibodies and S trimer was also higher in the SARS-CoV and SARS-CoV-2. In addition, we found that the ability of the HCoV-229E RBD to induce neutralizing antibodies was much lower and the intact and stable S1 subunit was essential for producing efficient neutralizing antibodies against HCoV-229E. Importantly, our results reveal different vaccine strategies for coronaviruses, and S-trimer is better than RBD as a target for vaccine development in *Alpha-coronavirus*. Our findings will provide important implications for future development of coronavirus vaccines.

**Importance:** Outbreak of coronaviruses, especially SARS-CoV-2, poses a serious threat to global public health. Development of vaccines to prevent the coronaviruses that can infect humans has always been a top priority. Coronavirus spike (S) protein is considered as a major target for vaccine development. Currently, structural studies have shown that *Alpha-coronavirus* (HCoV-229E) and *Beta-coronavirus* (SARS-CoV and SARS-CoV-2) RBDs are in lying and standing state, respectively. Here, we tested the ability of S-trimer and RBD to induce neutralizing antibodies among these coronaviruses. Our results showed that Beta-CoVs RBDs are in a standing state, and their S proteins can induce more neutralizing antibodies targeting RBD. However, HCoV-229E RBD is in a lying state, and its S protein induces a low level of neutralizing antibody targeting RBD. Our results indicate that *Alpha-coronavirus* is more conducive to escape host immune recognition, and also provide novel ideas for the development of vaccines targeting S protein.

## Introduction

Coronaviruses (CoVs) are enveloped, positive-sense, single-stranded RNA viruses with the largest genomes (26-32 kb) among known RNA viruses and are phylogenetically divided into four genera (*Alpha-, Beta-, Gamma*-, and *Delta-CoV*) (1, 2). To date, seven human-infecting coronaviruses (hCoVs) (3, 4) cause varying degrees of symptoms: HCoV-229E, HCoV-NL63, HCoV-HKU1, HCoV-OC43, severe acute respiratory syndrome coronavirus (SARS-CoV), Middle East respiratory syndrome coronavirus (MERS-CoV) and SARS-CoV-2, which is responsible for the current outbreak of COVID-19. Among them, alpha-CoVs (HCoV-229E and HCoV-NL63) and beta-CoVs (HCoV-OC43 and HCoV-HKU1) are well adapted to humans and widely circulate in the human population, with most infections causing mild disease in immunocompetent adults (3, 5, 6). In addition, SARS-CoV, SARS-CoV-2 and MERS-CoV belong to *Beta-CoV* and are highly pathogenic (7–9). SARS-CoV emerged in 2002 in the Guangdong Province of China and spread worldwide, resulting in 8,273 infections and nearly 775 deaths in 37 countries, with an approximately 9% case fatality rate (CFR) (7). MERS-CoV emerged in the Arabian Peninsula in 2012 and caused numerous outbreaks in humans, with a CFR of approximately 36% (10). SARS-CoV-2 is a new coronavirus strain that was first reported in Wuhan, China (4). As of June 5, 2020, SARS-CoV-2 has resulted in a total of 6,416,828 confirmed cases of COVID-19 worldwide and has caused 382,867 deaths (https://covid19.who.int/).

As the primary glycoprotein on the surface of the viral envelope, the spike (S) glycoprotein is the major target of neutralizing antibodies (nAbs) elicited by natural infection and key antigens in experimental vaccine candidates. The S protein contains two subunits responsible for receptor binding (S1 subunit) and membrane fusion (S2 subunit) (11). In particular, the S1 subunit of the prefusion S protein is structurally organized into four distinct domains: the N-terminal domain (NTD), the C-terminal domain (CTD), subdomain 1 (SD1) and subdomain 2 (SD2) (12–24).The receptor-binding domain (RBD) in the S protein mediates the binding of the virus to host cells, which is a critical step for the virus to enter target cells (11, 25). The CTDs of alpha-CoVs HCoV-NL63 and HCoV-229E are used as RBDs, which bind to angiotensin-converting enzyme 2 (ACE2) and aminopeptidase (APN), respectively (26, 27). The CTDs of beta-CoVs (SARS-CoV, SARS-CoV-2 and MERS-CoV) are similar in their core structures but are markedly different in their receptor-binding motifs (RBMs), leading to different receptor specificities; SARS-CoV and SARS-CoV-2 recognize ACE2 (28, 29), whereas MERS-CoV recognizes dipeptidyl peptidase-4 (DPP4) (30).

Cryo-electron microscopy (cryo-EM) studies have advanced our understanding of the S protein during virus entry. The S-trimer structures in the prefusion state have been reported for members of Alpha-CoVs (HCoV-NL63, HCoV-229E, porcine epidemic diarrhea virus (PEDV), and feline infectious peritonitis (FIPV) (12, 14, 16, 21), Beta-CoVs (mouse hepatitis virus (MHV), HCoV-HKU1, HCoV-OC43, SARS-CoV, SARS-CoV-2 and MERS-CoV) (13, 15, 19, 20, 22, 23), Gamma-CoV (avian coronavirus (IBV)) (24), and Delta-CoV (porcine deltacoronavirus (PDCoV)) (17, 18). The S1 subunits of Beta- and Gamma-CoV strains utilize the cross-subunit packing mode, reducing the conformational conflict of the RBD in a standing state (13, 19, 20, 24). In contrast, Alpha- and Delta-CoV strains both utilize an intrasubunit packing mode, and the S1-CTD is limited by the conformational conflict with surrounding domains (12, 14, 16–18, 21, 24). Hence, the S1-RBD in the S trimer was captured in two different states among different coronaviruses. In the Beta-CoVs (SARS-CoV, SARS-CoV-2 and MERS-CoV), the S1-RBD adopts a “standing” state, which is believed to be a prerequisite for receptor binding and RBM-specific antibody binding (13, 19, 20). Nevertheless, the S1-RBDs of alpha-CoVs all adopt “lying” state, which is considered more conducive to evading antibody recognition (12, 14, 16, 21).

Currently, no approved vaccines and drugs against the hCoVs are available. Past efforts, including the development of inactivated virus vaccines (31, 32), live-attenuated virus vaccines (33, 34), viral vector vaccines (35–37), subunit vaccines (38–40), DNA vaccines (41, 42), nanoparticle- (43) and virus-like particle (VLP)-based vaccines (44), were mainly focused on SARS-CoV and MERS-CoV. Compared with other vaccine types, subunit vaccines are target-specific and can generate high-titer nAbs without disadvantages, including viral infection, concerns of incomplete inactivation, virulence recovery and potential harmful immune responses. A number of subunit vaccines have been developed against SARS-CoV and MERS-CoV. Among them, the S protein or RBD was the major targets (45–47).

Compared with Beta-CoVs, relatively few studies have investigated two alpha-hCoVs: HCoV-229E and HCoV-NL63. However, their S1 subunit structure and receptor recognition pattern, especially the structure of the RBD and its state in the S trimer, differ substantially from those of beta-CoVs, suggesting different S protein immune responses between alpha- and beta-CoVs. Importantly, considering the low homology between different coronavirus genera, related research on alpha-CoVs can not only help to elucidate the differences between S proteins that adopt different RBD states but can also facilitate the development of coronavirus vaccines. In this study, we selected SARS-CoV, SARS-CoV-2, and HCoV-229E as models, which adopt the two RBD states, and evaluated and compared immune responses to the S trimers and RBDs of these coronaviruses through immunological and bioinformatics approaches. We also investigated the mechanism through which the HCoV-229E S trimer produced effective nAbs. Finally, we provide possible vaccine strategies for alpha- and beta-CoVs, which may facilitate the design and development of coronavirus vaccines in the future.

## Results and discussion

### Structural and immunological analyses of coronaviruses spike proteins

To date, many S-trimer structures of coronaviruses have been resolved (12–14, 16–24). Through structural comparison, we found an interesting phenomenon. Although the amino acid sequences are quite different, the compositions and structures of different functional domains of coronaviruses are similiar (**Fig. 1A**). All the alpha-CoVs bind to protein receptors with CTDs (as receptor binding domains (RBDs)), and the RBDs are in a lying state (12, 14, 16, 21, 26, 27) (**Fig.1B**). However, beta-CoVs (SARS-CoV, SARS-CoV-2 and MERS-CoV) bind receptors with CTDs (as RBDs), and the RBDs are in standing state (13, 19, 20) (**Fig. 1B**). Previous studies have shown that the RBD, as an important functional domain that directly binds to receptors, is an important target for the induction of nAbs (45–47). Hence, the transition between these two states (lying and standing) may play an important role in receptor binding and nAbs escape.

**Fig. 1.**
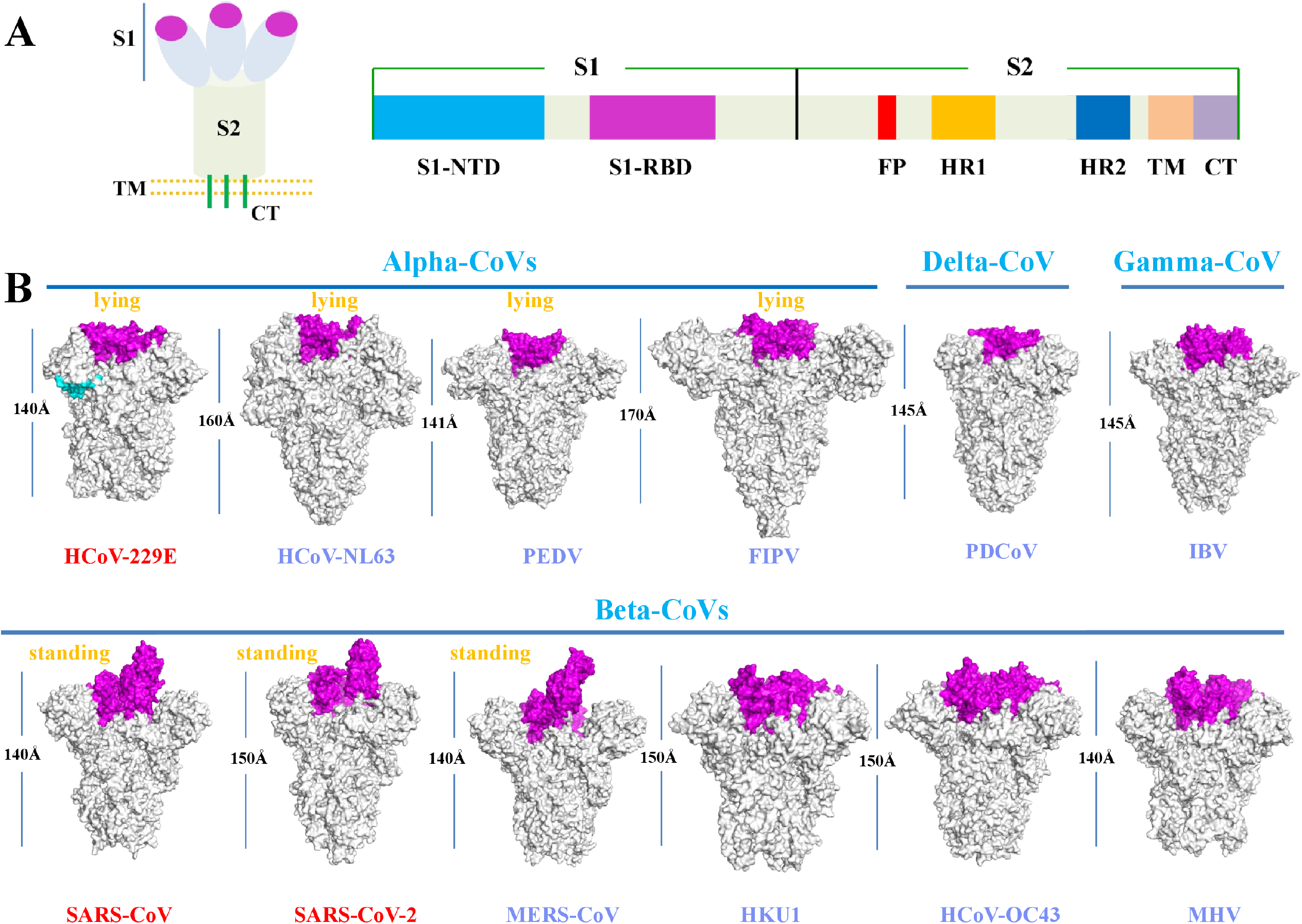
Structural analysis of S1-CTD from coronavirus S trimmers. (A) Schematic diagram of coronavirus spike protein organization. S1: receptor-binding subunit; S2: membrane fusion subunit; NTD: N-terminal domain; RBD: receptor-binding domain (magenta). (B) Overall structure comparison of coronavirus S trimers. The S trimer structures of HCoV-229E (PDB ID: 6U7H), HCoV-NL63 (PDB ID: 5SZS), PEDV (PDB ID: 6U7K), FIPV (PDB ID: 6JX7), PDCoV (PDB ID: 6BFU), IBV (PDB ID: 6CV0), SARS-CoV (PDB ID: 5X5B), SARS-CoV-2 (PDB ID: 6VSB), MERS-CoV (PDB ID: 5X5F), HKU1 (PDB ID: 5I08), HCoV-OC43 (PDB ID: 6OHW) and MHV (PDB ID: 3JCL) are shown. The S1-RBDs is colored in magenta. The lengths of the coronavirus structures are shown in previous reports.

To address this issue, we performed B-cell epitope predictions for the S trimers and RBDs of alpha-CoV (HCoV-229E) and beta-CoVs (SARS-CoV and SARS-CoV-2). The predicted positive residues (the corresponding spatial epitope and linear epitope) are displayed on the structural surface (**Fig. 2A, 2C and 2E**), and the distribution of positive residues on the RBD is summarized in **Table 1**. A total of 51 and 26 amino acid residues located on the RBD were predicted to be conformational epitopes for SARS-CoV and SARS-CoV-2, respectively. Of these, 47 and 25 residues were located in the SARS-CoV RBM subdomain and in the SARS-CoV-2 RBM subdomain, respectively. The linear B-cell epitope prediction results were similar in SARS-CoV and SARS-CoV-2. However, in HCoV-229E, only 3 residues located in the RBM subdomain were predicted to be conformational epitopes, and 9 residues were predicted to be linear epitopes. The same results also appeared in the HCoV-229E S trimer: fewer positive residues were located in the RBD than in the SARS-CoV or SARS-CoV-2 RBM subdomain (**Fig. 2A, 2C and 2E**). To better understand this result from a structural perspective, we analyzed the binding area of the RBD and receptors from SARS-CoV, SARS-CoV-2 and HCoV-229E (**Fig. 2B, 2D and 2F**). Among them, the interaction areas of SARS-CoV and SARS-CoV-2 were similar (approximately 829.7Å^2^ and 843.5Å^2^, respectively), which were much larger than that of HCoV-229E (approximately 497Å^2^). Furthermore, surface area analysis also yielded consistent results. Compared with HCoV-229E, the larger surface areas and binding areas of the SARS-CoV and SARS-CoV-2 RBDs to the receptor may induce more nAbs. Furthermore, the RBD of SARS-CoV and SARS-CoV-2 is in a standing state, the RBD in S-trimer can also induce higher levels of neutralizing antibodies than 229E.

**Fig. 2.**
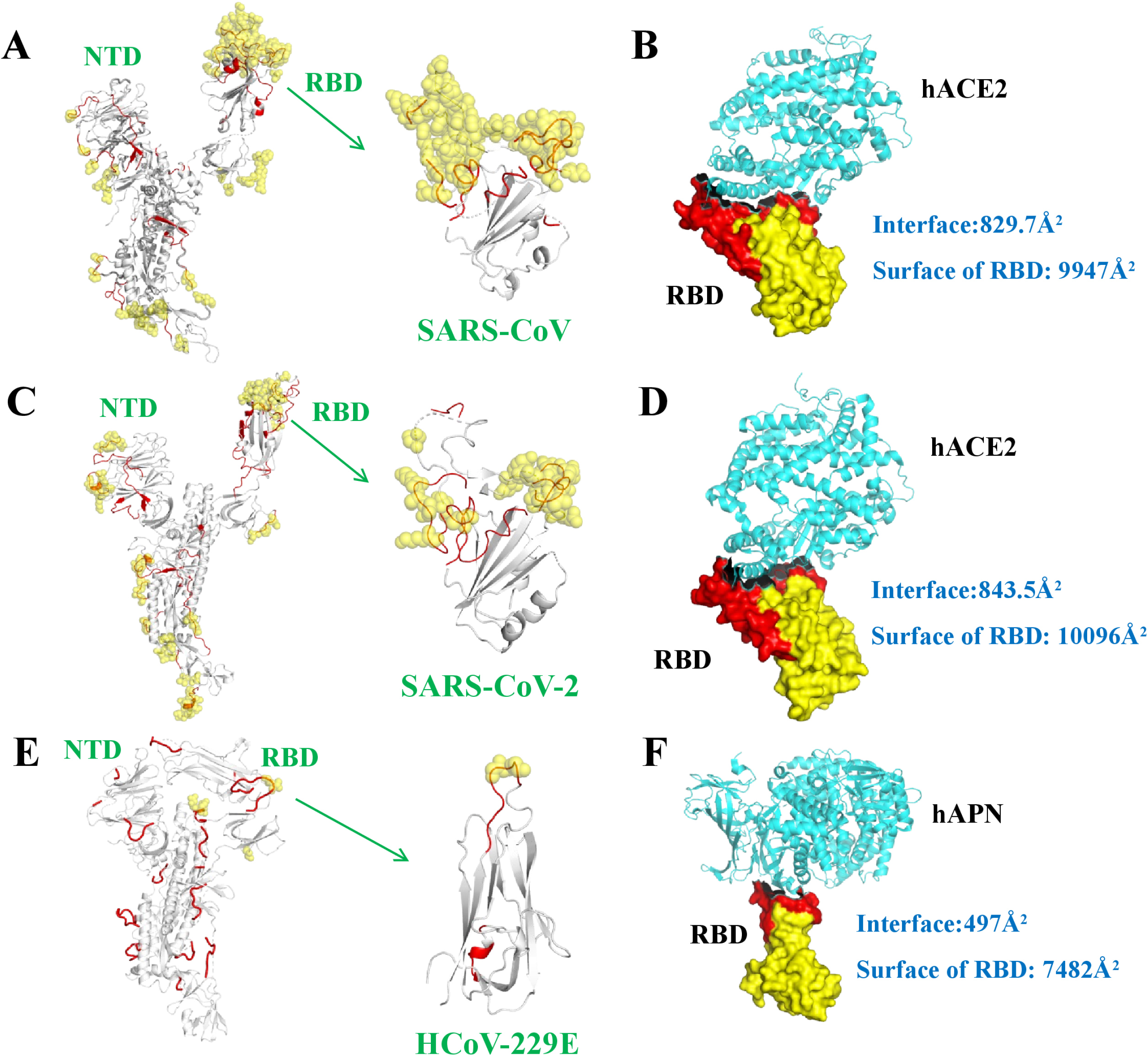
Structure-based B-cell epitope predictions of *Beta-CoV* (SARS-CoV and SARS-CoV-2) and *Alpha-CoV* (HCoV-229E). (A, C and E) The predicted B cell epitopes of SARS-CoV, SARS-CoV-2 and HCoV-229E are shown. The linear (red cartoon) and conformational (yellow sphere) B cell epitopes were predicted by Bepipred 2.0 or Discotope 2.0 and labeled onto the corresponding structure by PyMOL. (B, D and F) The complex structures of the RBDs of SARS-CoV, SARS-CoV-2 and HCoV-229E with the receptors (hACE2 and hAPN) are shown. The interface area of each complex and the surface area of each RBD were calculated via PDBePISA. The RBM region of the RBD and the receptors (hACE2 and hAPN) are shown in red and cyan, respectively.

**Table1.** Distribution of residues predicted positive for B cell epitopes

|  | RBD |  | RBM |  | RBM/RBD ratio |
| --- | --- | --- | --- | --- | --- |
|  | Conformational | Linear | Conformational | Linear | (%) |
| SARS-CoV | 51 | 56 | 47 | 33 | 92.2/58.9 |
| SARS-CoV-2 | 26 | 59 | 25 | 35 | 96.2/59.3 |
| HCoV-229E | 3 | 14 | 3 | 9 | 100/64.3 |
| HCoV-NL63 | 4 | 33 | 4 | 19 | 100/57.6 |
| PRCoV | 4 | 41 | 4 | 27 | 100/65.9 |
| TGEV | 7 | 34 | 4 | 24 | 100/70.6 |
| PEDV | 0 | 21 | 0 | 8 | 0/38.1 |
| FIPV | 0 | 17 | 0 | 11 | 0/64.7 |

### Distinct immunogenicity of the RBDs in *Alpha-CoV* (HCoV-229E) and *Beta-CoV* (SARS-CoV and SARS-CoV-2)

To evaluate the immunogenicity of the S trimers and RBDs between *Alpha-CoV* (HCoV-229E) and *Beta-CoV* (SARS-CoV and SARS-CoV-2), the S trimer and RBD were used as antigens to immunize mice. The antibody response was measured by ELISA using collected sera. The data showed that the S trimers and RBDs of SARS-CoV and SARS-CoV-2 could induce high levels of protein-specific antibodies (antibody titers: 1.36×10^5^, 3.84×10^5^; 1.625×10^6^, 5×10^5^, respectively, **Fig. 3A–3D**). Moreover, the S trimers of both SARS-CoV and SARS-CoV-2 could induce high-titer RBD antibodies (antibody titers: 1.28×10^5^; 2.75×10^5^, respectively), and the RBD-specific antibodies had a high affinity for the S trimer (antibody titers: 6.72×10^5^; 5×10^5^, respectively, **Fig. 3A–3D**). Similar to SARS-CoV and SARS-CoV-2, the S trimer and RBD of HCoV-229E both had good immunogenicity (antibody titers: 8.8×10^4^; 7.68×10^5^, respectively, **Fig. 3E and 3F**). However, the HCoV-229E S trimer induced fewer RBD-specific antibodies than those of SARS-CoV and SARS-CoV-2 (antibody titer: <500, **Fig. 3F**), and the HCoV-229E RBD-specific antibodies had a lower affinity for the S trimer (antibody titer: 1.125×10^3^, **Fig. 3E**). To confirm this finding, a higher immunization dose of the HCoV-229E RBD (50 μg) was used in the same manner. Nevertheless, no significant increase in the RBD-specific antibody titer (antibody titers: 1.792×10^6^, 3.072×10^6^, **Fig. 3G**) or antibody affinity for the HCoV-229E S trimer (antibody titers: 2.5×10^2^, 2.5×10^2^, **Fig. 3H**) was noted. Overall, our immune epitope analysis and biochemical tests consistently showed that the S trimer with a “standing” RBD state that is more conducive to inducing RBD-specific antibodies in SARS-CoV and SARS-CoV-2. The “lying” RBD state induces fewer RBD-specific antibodies and resulted in a lower affinity between the S trimer and the RBD-specific antibodies in HCoV-229E.

**Fig. 3.**
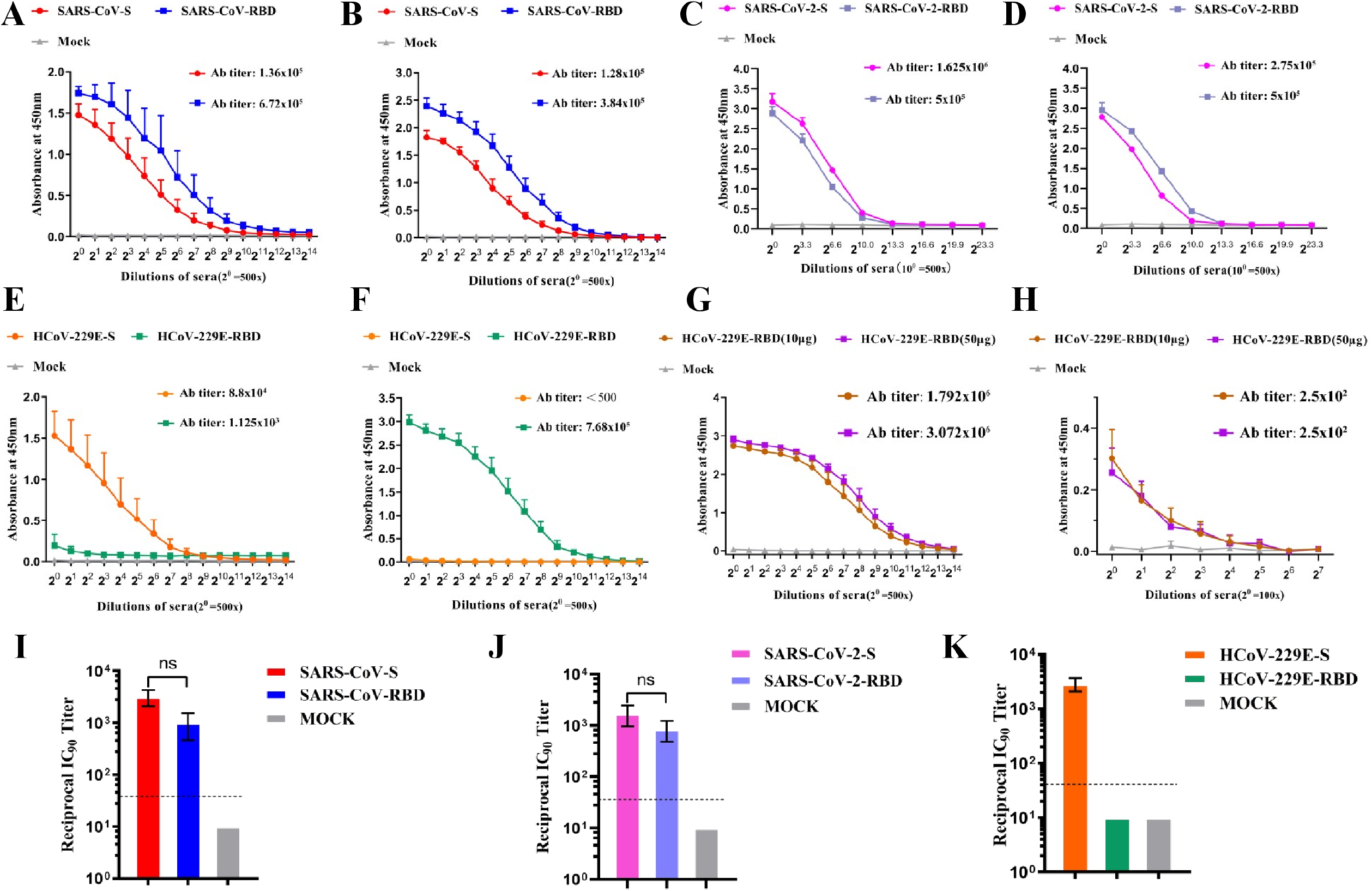
Immunological analysis of *Beta-CoV* (SARS-CoV and SARS-CoV-2) and *Alpha-CoV* (HCoV-229E). (A and B) Cross-reactivity of the SARS-CoV S trimer and RBD-specific sera is determined by ELISA. Mice sera of SARS-CoV S trimer (red) and SARS-CoV RBD (blue) were 10-fold serially diluted (starting with 500-fold dilution) and reacted with the S trimer (A) or RBD (B), respectively. (C and D) Cross-reactivity of the SARS-CoV-2 S trimer and RBD-specific sera is determined by ELISA. Mice sera of SARS-CoV-2 S trimer (magenta) and SARS-CoV-2 RBD (slate) were 2-fold diluted and reacted with SARS-CoV-2 S trimer (C) and RBD (D). (E and F) Cross-reactivity of the HCoV-229E S trimer and RBD-specific sera is determined by ELISA. Mice sera of HCoV-229E S trimer (orange) and HCoV-229E RBD (green) were 2-fold diluted and reacted with HCoV-229E S trimer (E) and RBD (F). (G and H) The antibody titers of sera from mice immunized with 10 μg of the HCoV-229E RBD (brown) and 50 μg of the HCoV-229E RBD (purple). Mice sera were reacted with the HCoV-229E RBD (G) or the spike trimer (H). All data above are presented as the mean A450 ± s.e.m and the IgG antibody titers of each serum were calculated as the maximum endpoint dilution that remained positive. (I, J and K) The neutralization assay of mouse sera from the spike trimer and RBD against SARS-CoV, SARS-CoV-2 and HCoV-229E pseudoviruses is determined. The data are presented as the mean reciprocal IC_90_ titer. The limit of detection for the assay depends on the initial dilution and is represented by dotted lines, a reciprocal IC_90_ titer of 10 was assigned.

### *Alpha-CoV* (HCoV-229E) induced fewer RBD-specific neutralizing antibodies

Next, we tested the neutralizing ability of the sera using a vesicular stomatitis virus (VSV)-based pseudovirus. Both the S trimer and RBD sera from SARS-CoV- and SARS-CoV-2-immunized mice had a good neutralizing ability (**Fig. 3I and 3J**). For HCoV-229E, the S trimer serum had a comparable neutralizing ability to that of SARS-CoV or SARS-CoV-2, but the RBD serum had no detectable neutralizing ability (**Fig. 3K**). Our experimental results indicate that the lying state of the RBD in the HCoV-229E S-trimer induces the production of very few antibodies targeting the RBD, but the S-trimer still produces strong neutralizing antibody levels.

In this study, we found that more RBD-specific antibodies were induced by the S trimer with the RBD in the standing state than the S trimer with the RBD in the lying state, and the affinity between RBD-specific antibodies and the S trimer was also higher in the standing state. However, we also found that fewer nAbs were induced by the RBD of HCoV-229E than by the RBDs of SARS-CoV or SARS-CoV-2. In terms of HCoV-229E, the distribution of the potential residues in the RBM was lower than that of SARS-CoV or SARS-CoV-2, which may have been caused by different RBM patterns and exposure degrees. When we compared the reported nAb epitopes of SARS-CoV and *Alpha-CoV* TGEV with our results (47), they were basically consistent. Therefore, we believe that this finding illustrates the inherent difference between the RBDs of *alpha*- and *beta-CoV*.

### The intact and stable S1 subunit of HCoV-229E is a prerequisite for the production of effective nAbs

Our experimental results showed that HCoV-229E S-trimer can induce strong nAb levels, while the RBD alone is less immunogenic. Next, we will explore which functional domains of the S-trimer are involved in the generation of nAbs. To clarify this issue, we immunized mice with the HCoV-229E S trimer (10 μg), S1 (10 μg), NTD (10 μg), RBD (10 μg) and NTD+RBD (5 μg+ 5 μg). Meanwhile, to better confirm our results, the HCoV-229E strain VR740 was used for the neutralizing assay. The results indicated that the S trimer serum had the best neutralizing ability, followed by the S1 and NTD+RBD sera, while the NTD and RBD sera alone had no detectable neutralizing effects (**Fig. 4A**). The results indicate that the S1 region in the S-trimer should be the key region for nAbs induction. To further verify the importance of the complete S1 structure in the S-trimer, we designed two S trimer mutants, namely, an NTD-deficient S trimer and an S65C/T472C S trimer, the S1 subunit integrity or stability of which was destroyed (**Fig. 4C and 4F**). Mutant proteins disrupt the conformational conflicts that limit RBD standing, significantly improving their ability to bind hAPN (**Fig. 4D and 4G**). However, an incomplete or unstable S1 conformation significantly reduces the level of nAbs induced by the S-trimer (**Fig. 4E and 4K**). Taken together, these results showed that the intact and stable S1 subunit of HCoV-229E is a prerequisite for the production of effective nAbs.

**Fig. 4.**
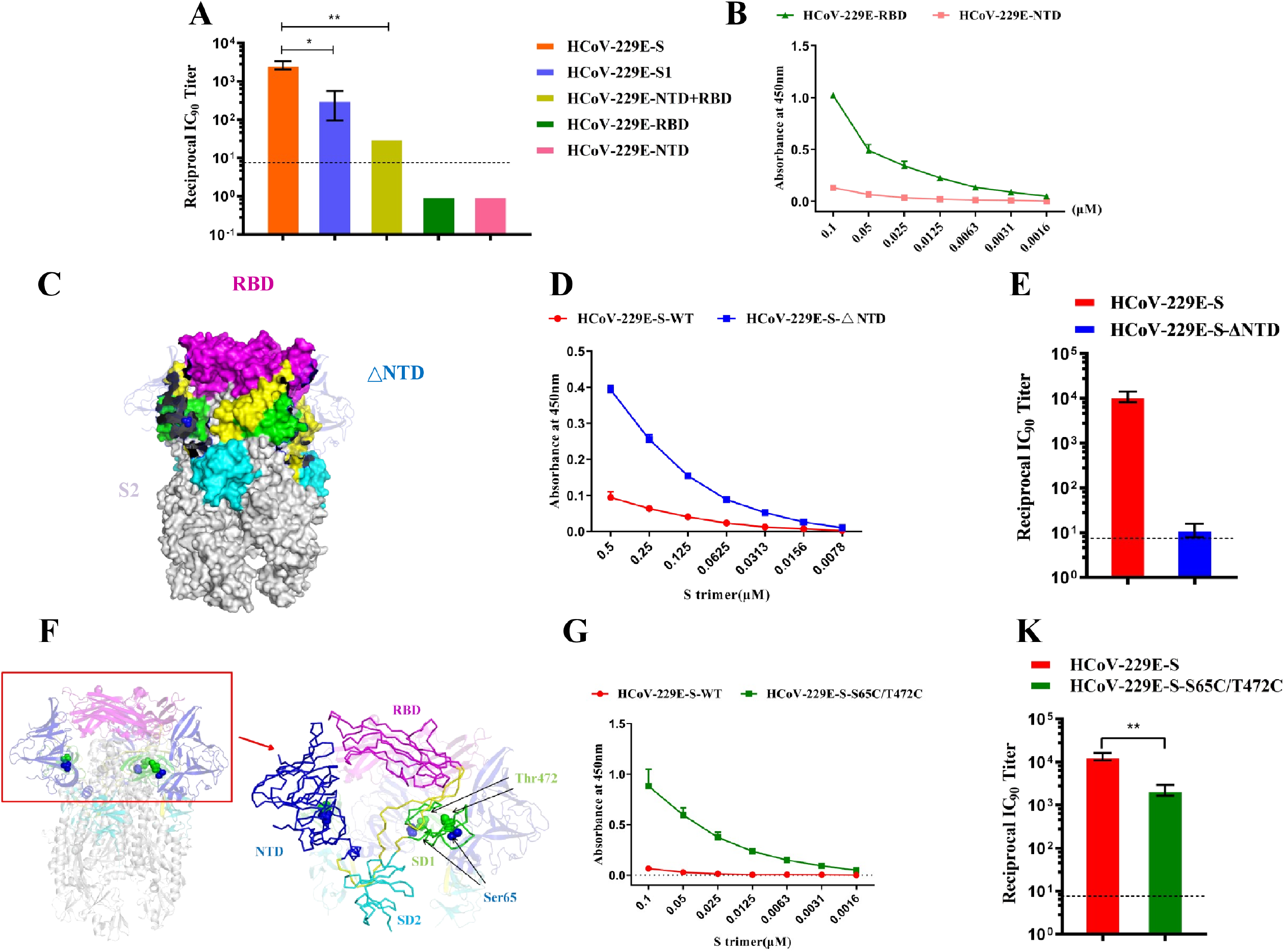
The intact and stable S1 subunit of HCoV-229E is a prerequisite for the production of effective nAbs. (A) The neutralization abilities of mouse sera from the HCoV-229E S trimer, S1, NTD+RBD, NTD and RBD against HCoV-229E strain VR740. (B) Determination of the affinity of NTD and RBD with the receptor hAPN. (C) Structural model of HCoV-229E-S-ΔNTD. Magenta: RBD; green: SD1; cyan: SD2. (D) Dose-dependent binding of HCoV-229E-S-ΔNTD and hAPN. (E) The neutralization ability of mouse sera from HCoV-229E-S-ΔNTD was measured via pseudovirus neutralization assay. (F) The structure of HCoV-229E-S-S65C/T472C. Ser65 and Thr472 are shown in spheres in the magnified region. Magenta: RBD; blue: NTD; green: SD1; cyan: SD2. (G) Dose-dependent binding of HCoV-229E-S-S65C/T472C and hAPN. (H) The neutralization ability of mouse sera from HCoV-229E-S-S65C/T472C was measured via pseudovirus neutralization assay. In the neutralization assay, the data are presented as the mean reciprocal IC_90_ titer (n=4). The limit of detection for the assay depends on the initial dilution and is represented by dotted lines, a reciprocal IC_90_ titer of 10 was assigned. Besides, data are presented as the mean OD_450_ ± s.e.m. (n=3) in ELISA assay.

Furthermore, our experimental results show that RBD has a higher ability to bind to the receptor hAPN (**Fig. 4B**), which indicates that the characteristics of RBD itself may lead to the generation of less neutralizing antibodies. Furthermore, we screened monoclonal antibodies using S-trimer, and the results showed that few antibodies targeting S1-RBD (**Fig. 5A**). To further determine the ability of RBD to induce antibodies itself, we screened monoclonal antibodies targeting the S1 region and found that the proportion of antibodies targeting RBD was approximately 20% (**Fig. 5B**). Since the S1 protein is expressed in a monomeric form, RBD is not restricted by the conformation of the surrounding domains and should be in a standing state. Therefore, our results indicate that HCoV-229E RBD may induce weaker levels of neutralizing antibodies, which may be related to its own characteristics. Furthermore, the RBD in the HCoV-229E S-trimer is in the lying state. Its characteristics and conformational state may prompt the HCoV-229E to escape the host’s immune surveillance, thereby allowing itself to circulate in the population for a long time.

**Fig. 5.**
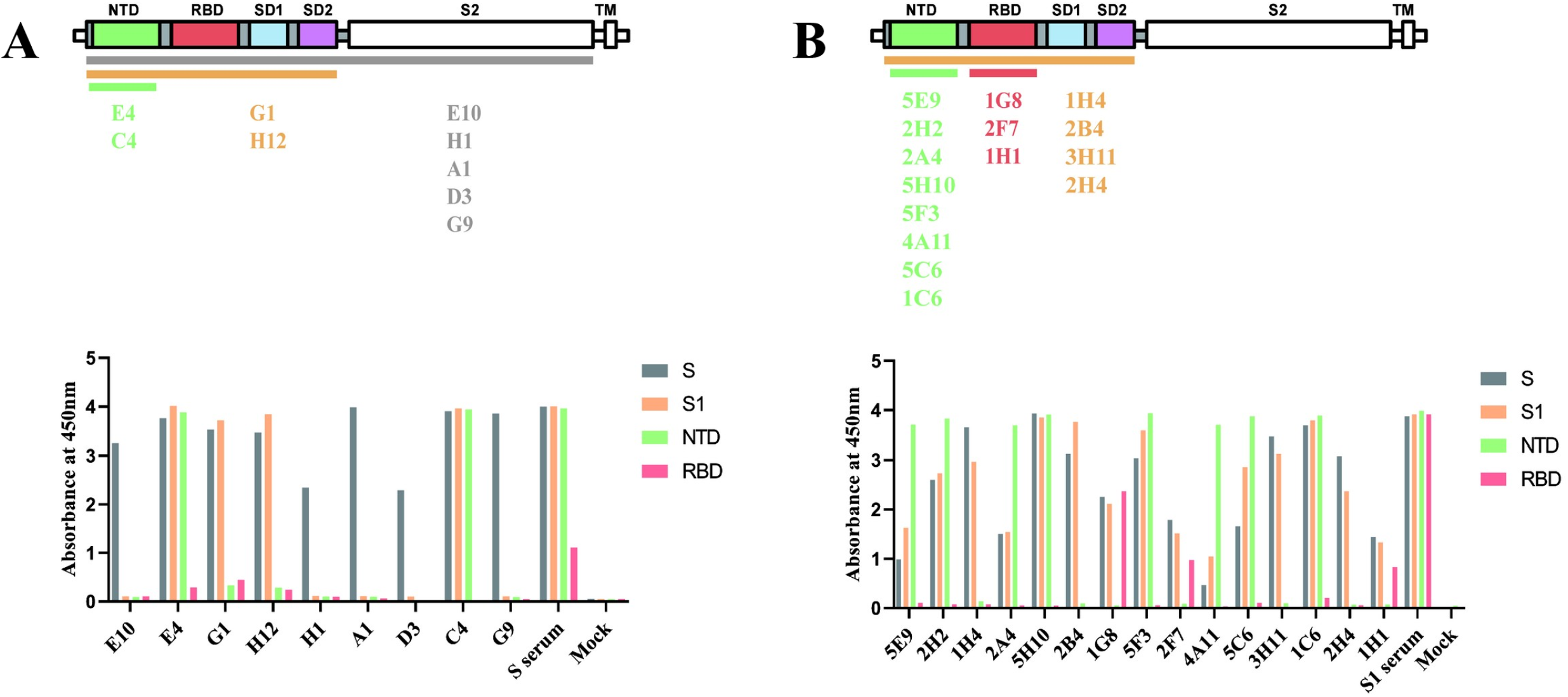
Monoclonal antibody epitope mapping of the HCoV-229E spike protein. Monoclonal antibody (MAb) epitope regions in the HCoV-229E spike protein (A) and S1 domain (B). Supernatants of positive hybridomas were reacted with the HCoV-229E spike protein, S1, NTD and RBD. Data are presented as the OD_450_ (bottom). MAbs and their epitope regions are indicated below the schematic of the HCoV-229E spike.

### Potential vaccine strategies for *Alphacoronavirus* (HCoV-229E) and *Betacoronavirus* (SARS-CoV and SARS-CoV-2)

We compared the structures of S trimers and RBDs among alpha-coronaviruses (**Figs. 1B and 6A**). We also predicted the potential B-cell epitopes for their RBDs (**Fig. 6A; Table1**). In *Alpha-CoV*, the S-trimer had a closed S1 subunit with three “lying” RBDs **(Fig. 1B)**. Moreover, the RBDs consist of a standard β-sandwich fold core and three short discontinuous loops in the same spatial region (12, 14, 16, 21, 26, 27, 48) (**Fig. 6A**). Meanwhile, we performed a structural conservative analysis and the results showed that the RBD structures of HCoV-NL63, PEDV, and FIPV are most similar to HCoV-229E, with RSMD values of 1.9, 2.0, and 2.2, respectively (**Fig. 6B**). In addition, the distribution of potential B-cell epitopes in the RBDs of alpha-CoVs was also similar to that of HCoV-229E (**Fig. 6A and 6C; Table1**). Based on the above data, inherent differences exist in the RBDs between alpha- and beta-CoVs (**Figs. 2 and 6A**). However, the alpha- and beta-CoVs show high similarity in their RBDs and similar potential immune characteristics within their respective genera (**Figs. 2, 3, 6A and 6B**). Accordingly, in alpha-CoVs such as HCoV-229E, subunit vaccines should prioritize the S-trimer rather than the RBD. In beta-CoVs such as SARS-CoV and SARS-CoV-2, the S trimer and RBD are both good candidates for subunit vaccines (**Fig. 7**).

**Fig. 6.**
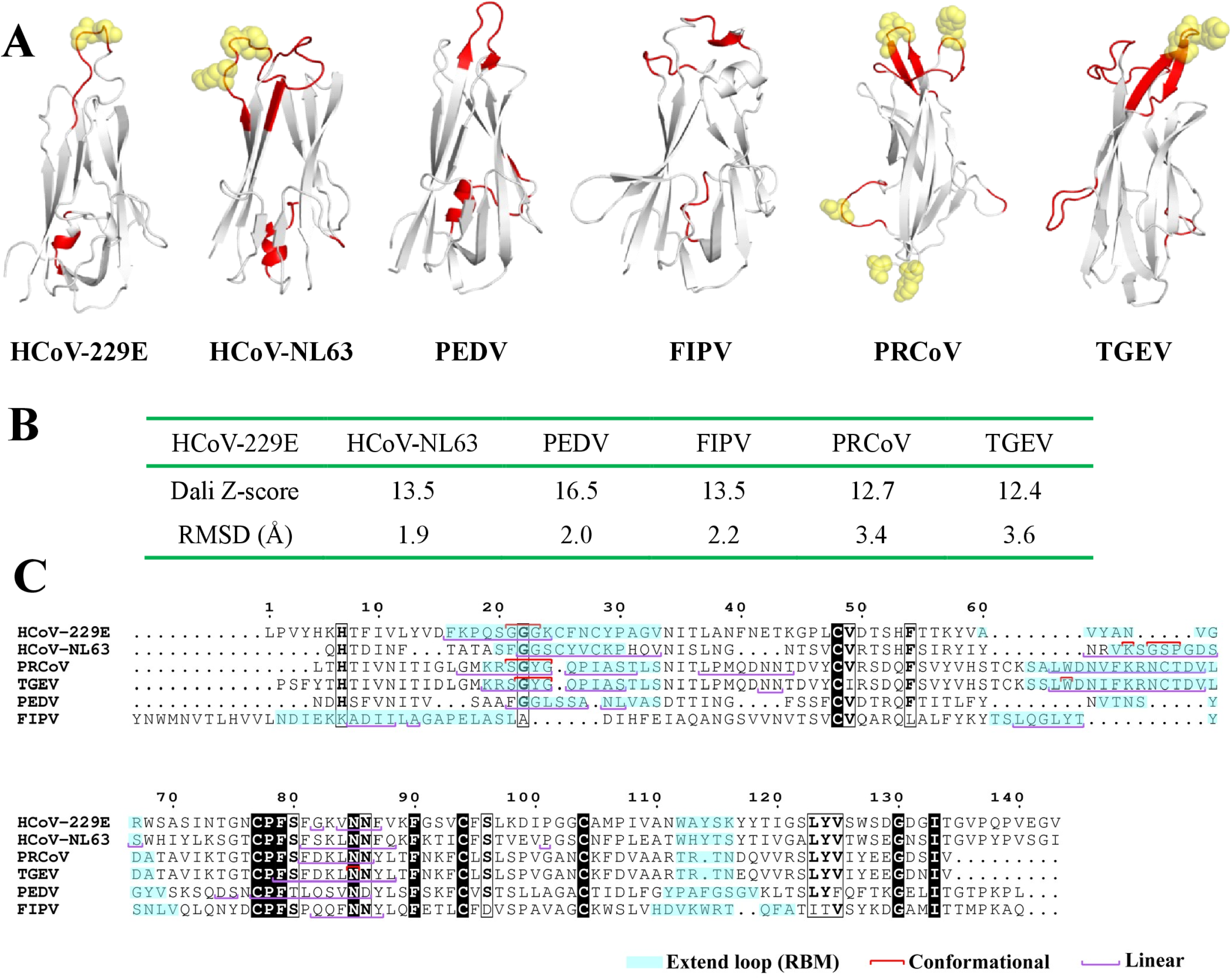
B cell epitope analysis of the RBD regions of *alpha-coronavirus* spike proteins. (A) Structures of the RBDs from alpha-CoVs (HCoV-229E, HCoV-NL63, PEDV, FIPV, PRCoV and TGEV) spike proteins. The linear (red cartoon) and conformational (yellow sphere) B cell epitopes were predicted by Bepipred 2.0 or Discotope 2.0 and labeled onto the corresponding RBD structure by PyMOL. (B) Structural comparison of the RBDs from alpha-CoVs. (C) Sequence alignment of the RBDs from alpha-CoVs. The RBM or putative RBM region is shown in cyan. The amino acid residues predicted for linear (purple) and conformational (red) B cell epitopes are also shown.

**Fig. 7.**
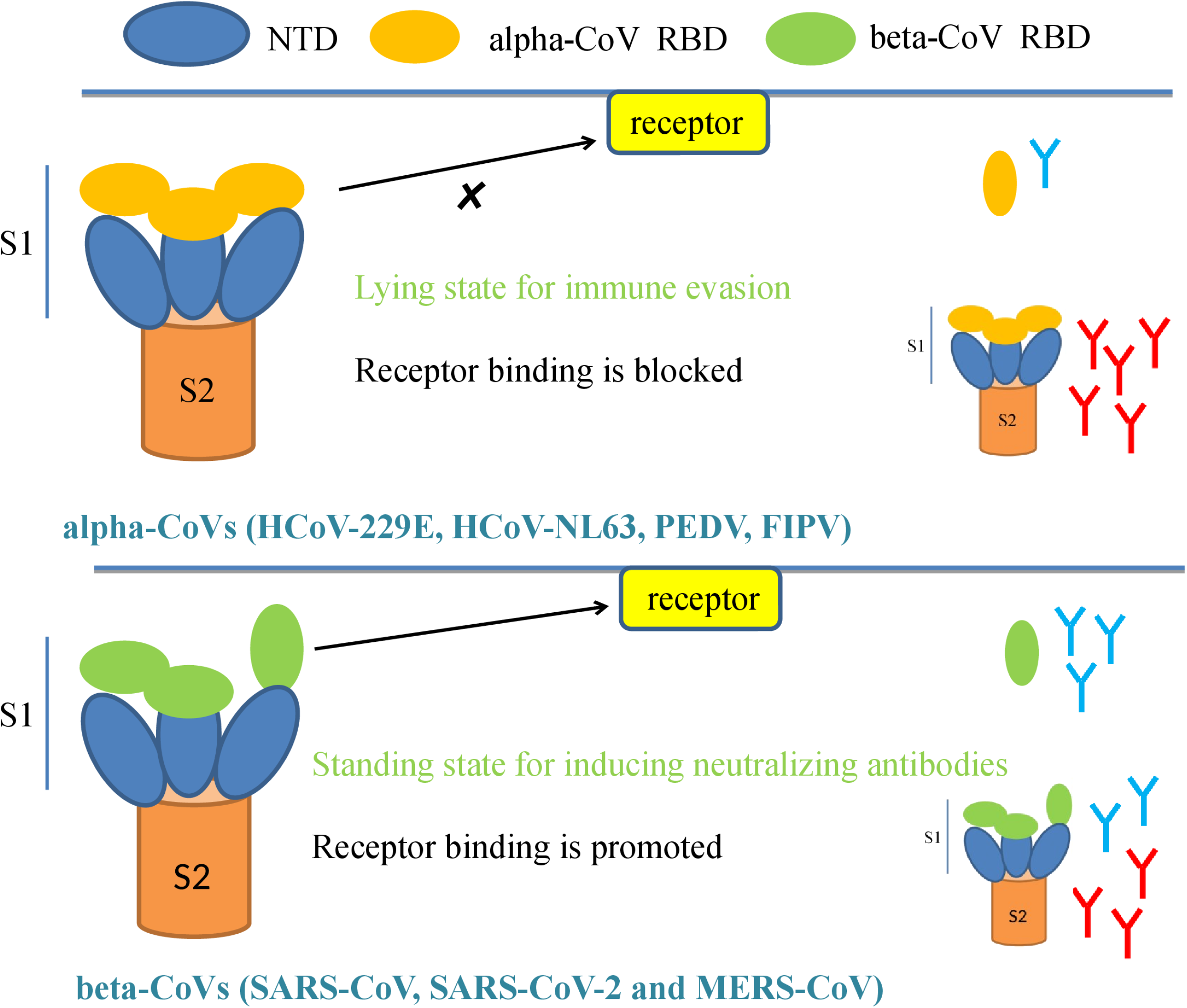
Potential vaccine strategies for alpha- and beta-CoVs. The model showed that the RBDs of the *alpha-CoV* S trimers are in a lying state. In this state, the S protein cannot bind to the receptor, but meanwhile, this state is also conducive to escaping the immune response target the RBD, and the RBDs of the alpha-CoVs also induces fewer NAbs; thus, their S-trimers can be an effective potential subunit vaccine. In beta-CoVs (SARS-CoV, SARS-CoV-2 and MERS-CoV), the RBDs of the S trimer are in a standing state, which is conducive to binding receptors, and the RBD can induce more antibodies; thus, their S-trimers and RBDs can produce more NAbs. Hence, their S-trimers and RBDs can be an effective potential subunit vaccine.

In summary, we systematically analyzed the conformational states and immunogenicity of the S-trimers and RBDs of *Alpha-CoV* (HCoV-229E) and *Beta-CoV* (SARS-CoV and SARS-CoV-2). Our results showed that the inherent differences between the RBDs of alpha- and Beta-CoVs and revealed potential identical immune characteristics in alpha- and Beta-CoVs. Based on these findings, we provide potential vaccine strategies for alpha- and Beta-CoVs: for alpha-CoVs, the S trimer or S1 subunit is more suitable for subunit vaccines than the RBD, but the ADE effect of the alpha-CoVs S trimer still requires further investigation. For Beta-CoVs, SARS-CoV and SARS-CoV-2, the S trimer and RBD are both candidates for subunit vaccines. However, considering the ADE effect reported in SARS-CoV and the homology between the SARS-CoV and SARS-CoV-2 S proteins, the RBD may be a priority in the design of subunit vaccines. Although our inference requires more experimental data for further confirmation, our results will provide a reference for the development of coronavirus vaccines in the future.

## Materials and methods

### Plasmid construction

According to previous research, insect codon-optimized sequences encoding the HCoV-229E S glycoprotein ectodomain (GenBank accession number NP_073551.1, residues 1-1,116) and SARS-CoV S glycoprotein ectodomain with an R667A mutation (GenBank accession number NP_828851.1, residues 1-1,195) were cloned into the baculovirus transfer vector pFastbac1 (Invitrogen) with a gene fragment encoding the GCN4 trimerization motif (LIKRMKQIEDKIEEIESKQKKIENEIARIKKIK) and an eight-residue Strep-tag (WSHPQFEK) (20, 22). Additionally, human aminopeptidase N (hAPN) (GenBank accession number JX869059, residues D66-K967) (49) was cloned into the pFast-bac1 vector with an N-terminal honeybee melittin signal peptide and a C-terminal 6x His-tag. HCoV-229E S1 (M1-A536), S1-NTD (M1-V258) and S1-RBD (295V-428V) containing an N-terminal honeybee melittin signal peptide and a C-terminal Fc-tag were constructed using the same method (29). Besides, the S fragments of HCoV-229E, SARS-CoV and SARS-CoV-2 were cloned into the pcDNA3.1 (+) vector with a C-terminal His-tag using a previously described protocol (50). All constructs were validated by DNA sequencing. The S protein sequences of HCoV-229E, SARS-CoV and SARS-CoV-2 (GenBank accession number NC_045512.2) were synthesized by GenScript Corporation (GenScript, Nanjing, China).

### Protein expression and purification

The spike protein ectodomain (including a variety of truncated proteins and mutant proteins) and hAPN were expressed and purified using a previously described protocol (20, 29). Briefly, the construct was transformed into bacterial DH10Bac competent cells (Invitrogen); then, the extracted bacmid was transfected into Sf9 cells (American Type Culture Collection). The supernatant of the cell culture containing the secreted S glycoprotein was harvested at 60 h after infection and concentrated, and the buffer was changed to binding buffer (10 mM HEPES pH 7.2 and 500 mM NaCl). Finally, the S glycoprotein was captured by StrepTactin Sepharose High Performance resin (GE Healthcare) and eluted with 10 mM D-desthiobiotin in the binding buffer (20). For SARS-CoV-2, the S-trimer ectodomain and RBD were purchased from Sino Biological, Inc. Finally, the protein storage buffer is exchanged for 10 mM HEPES pH 7.2 and 150 mM NaCl for subsequent assays.

### Animal immunization

Female BALB/c mice aged 6 weeks were immunized with different proteins at 0 and 3 weeks. Proteins (10 μg) diluted in HEPES-buffered saline (HBS; 10 mM HEPES and 150 mM NaCl) were mixed 1:1 with the 2× Sigma Adjuvant System. The mice were intramuscularly inoculated with 50 μl of this solution (25 μl into each hind leg). Two weeks after the final immunization, sera were collected for subsequent assays. For SARS-CoV-2, the corresponding rabbit polyclonal antibodies (pAbs) were purchased from Sino Biological, Inc.

### Enzyme-linked immunosorbent assay (ELISA)

To measure the immune responses of different sera, ELISA plates were coated with purified protein at 0.1 μM/well in citrate-buffered saline (CBS, pH=9.6) overnight at 4°C and subsequently blocked with phosphate-buffered saline (PBS) with 0.05% Tween 20 (PBST) containing 1% bovine serum albumin (BSA, w/v) at 37°C. After standard washes, the plates were incubated with 2- or 10-fold serially diluted sera for 1 h at 37°C. Then, horseradish peroxidase (HRP)-conjugated goat anti-mouse/rabbit IgG (1:10,000 diluted in PBST with 1% BSA (w/v), Boster) was used as the secondary antibody, and 3,3’,5,5’-tetramethylbenzidine (TMB) (Beyotime) was used as the substrate for detection. Optical density (OD) was read at 450 nm and 630 nm using a SPARK10M microplate reader (TECAN) after stopping the reaction with 2 M H_2_SO_4_. Sera from mice immunized with HBS were used as a mock control.

For the receptor-binding assay, the ELISA plates were coated with hAPN at 0.1 μM/well in CBS (pH=9.6) overnight at 4°C and subsequently blocked with PBST containing 1% BSA (w/v) at 37°C. After washing, the HCoV-229E S-trimer, NTD, RBD and mutant proteins were serially diluted 2-fold in HBS and incubated with the plates for 1h at room temperature. Then, the mouse anti-Strep-tag II antibody (SAB, 1:3,000 diluted in PBST with 1% BSA (w/v)) and HRP-conjugated goat anti-mouse IgG (1:5,000 diluted in PBST with 1% BSA (w/v), Boster) was used for detection. Signal reading was carried out in the same manner. HBS buffer was used as a mock control.

### Generation of HCoV-229E mAbs and epitope mapping

Six-week-old female BALB/c mice were immunized with 100 μg of purified HCoV-229E S-trimer or S1 protein. Antigens were emulsified in Freund’s Complete Adjuvant (Sigma-Aldrich, F5881) for the first immunization or Freund’s Incomplete Adjuvant (Sigma-Aldrich, F5506) for the subsequent boost. Each mouse received three subcutaneous injections at two-weeks intervals. Mice with the highest titers of antibodies against the HCoV-229E S-trimer or S1 protein were further boosted by intraperitoneal injection 200 μg of purified HCoV-229E S-trimer or S1 protein diluted in PBS buffer. Three days after the last injection, spleen cells were collected and fused with SP2/0 cells with PEG1450 (Sigma-Aldrich, P7181) to generate hybridoma cells. Antigen-specific ELISA was used for the hybridoma screening. Positive hybridomas were further subcloned and used for epitope mapping. To this end, ELISA plates were coated with different proteins (the HCoV-229E S-trimer, S1, NTD and RBD) at 1μg/ml in CBS (pH=9.6) overnight at 4°C and subsequently blocked and washed. Then the plates were reacted with the hybridoma culture supernatants at 37°C for 1h. HRP-conjugated goat anti-mouse IgG (1:5,000 diluted in PBST with 1% BSA (w/v), Boster) was used for detection. Signal reading was carried out in the manner described above. Hybridoma culturing medium was used as a mock control.

### Production and entry assay of pseudoviruses

Pseudo-typed viruses were produced as previously described (50), 293T (ATCC, CRL-3216), Huh-7 and Vero (ATCC, CCL-81) cells were maintained in high glucose DMEM (Gibco, USA) supplemented with 10% FBS (FBS; Natocor, Argentina), penicillin (100 IU/ml) and streptomycin (100 μg/ml). Human coronavirus 229E (ATCC, VR-740™) was amplified by Huh-7 cells.

The 293T cells (T25) were transfected with 1 μg of HCoV-229E-S-Δ19-pcDNA3.1 (C-terminal deletion 19aa), SARS-CoV-S-Δ22-pcDNA3.1 plasmids (C-terminal deletion 22 aa) and SARS-CoV-2-S-Δ18-pcDNA3.1 plasmids (C-terminal deletion 18 aa). Additionally, VSV-G-pcDNA3.1 and pcDNA3.1 were transfected as positive and negative controls, respectively, using Exfect2000 transfection reagent (Vazyme). Twenty-four hours later, the transfected cells were infected with VSV-ΔG-G at 1 MOI. Twenty-four hours post infection, VSV-ΔG-HCoV-229E-S, VSV-ΔG-SARS-CoV-S and VSV-ΔG-SARS-CoV-2-S (culture supernatants) were harvested and centrifuged at 10,000 rpm/min for 10 min, and the supernatant was collected and stored at −80°C in 2ml aliquots until use.

For titration of these three pseudovirus, 2-fold dilution was performed in hexaplicate wells of 96-well culture plates. The last column served as the cell control with added pcDNA3.1-transfected supernatant. After 48 h of incubation in a 5% CO_2_ environment at 37°C, the culture supernatant was removed and washed by PBS three times, and 20 μl of Reporter Lysis 5X Buffer was added to each well to complete a single freeze-thaw cycle (Promega, Cat. # E4030). Then, the luciferase substrate (Promega, Cat. #E1500) was added to each well for luminescence detection using a Multimode Microplate Reader (Tecan Spark 10M).

For pseudovirus neutralization experiments, 2-fold serial dilutions of mouse sera (initially 1:40) were mixed with the three pseudovirus strains, which were previously titered to target approximately 50,000 RLU. After 48 h of incubation, the RLU value was read. According to the formula ((1- (x-c)/x)%; x: sample reading, c: cell control reading, n=3), the neutralization protection rate was calculated. For virus neutralization experiments, 2-fold serial dilutions of mouse sera (initially 1:8) were mixed with an equal volume of H229E virus (100 TCID50 /well) at 37°C. The neutralization titers were measured by the observed CPE. Serum from a PBS-treated mouse was used as a negative control.

### B-cell epitope prediction analysis

According to previous research, B-cell epitope were predicted and analyzed (IEDB, http://www.iedb.org) (51). Briefly, structure-based B-cell epitope prediction was performed by DiscoTope 2.0 with a positive cutoff greater than −3.7 (corresponding to a specificity greater than or equal to 0.75 and a sensitivity less than 0.47) using the following protein structures: the HCoV-229E S-trimer and RBD (PDB IDs: 6U7H and 6ATK, respectively), the SARS-CoV S-trimer and RBD (PDB IDs: 5X5B and 2AJF, respectively), the SARS-CoV-2 S-trimer and RBD (PDB IDs: 6VYB and 6M0J, respectively), the PEDV RBD (PDB ID: 6U7K), the FIPV RBD (PDB ID: 6JX7), the PRCoV RBD (PDB ID: 4F5C), and the transmissible gastroenteritis virus (TGEV) RBD (PDB ID: 4F2M). For linear B-cell epitope prediction, the corresponding amino acid sequences from the above structures were used. The BepiPred 2.0 algorithm was applied with a cutoff of 0.55 (corresponding to a specificity greater than 0.81 and a sensitivity less than 0.3). All the predicted residues were then labeled in corresponding structures using PyMOL (Schrödinger). The interaction area and surface area were analyzed via PDBePISA (https://www.ebi.ac.uk/msd-srv/prot_int/pistart.html). Additionally, the amino acid sequences of alpha-CoVs RBDs were aligned using ClustalW2 (52). The NCBI accession numbers of the sequences used were as follows: HCoV-229E (AAQ90002.1), HCoV-NL63 (AVL25587.1), PRCoV (AAA46905.1), TGEV (CAB91145.1), PEDV (AIU98611.1) and FIPV (ACT10887.1).

### Statistical analysis

Statistical significance was determined using an unpaired two-tailed Student’s t test. Values <0.05 were considered statistically significant. All experiments were further confirmed using biological repeats.

### Ethics statement

All the mice used in this study were maintained in compliance with the recommendations in the Regulations for the Administration of Affairs Concerning Experimental Animals established by the Ministry of Science and Technology of China. The experiments were carried out using the protocols approved by the Scientific Ethics Committee of Huazhong Agricultural University (permit number: HZAUSW-2018-009).

## Acknowledgments

This work was supported by National Natural Science Foundation of China Grants 31722056 and 31702249, National Key R&D Plan of China Grant 2018YFD0500100, China Postdoctoral Science Foundation Grant 2019M662674 and the Huazhong Agricultural University Scientific and Technological Self-innovation Foundation (program no. 2662017PY028).

The authors declare no competing interests.

